# Vaccine-elicited murine antibody WS6 neutralizes diverse beta-coronaviruses by recognizing a helical stem supersite of vulnerability

**DOI:** 10.1101/2022.01.25.477770

**Authors:** Wei Shi, Lingshu Wang, Tongqing Zhou, Mallika Sastry, Eun Sung Yang, Yi Zhang, Man Chen, Xuejun Chen, Misook Choe, Adrian Creanga, Kwan Leung, Adam S. Olia, Amarendra Pegu, Reda Rawi, Chen-Hsiang Shen, Erik-Stephane D. Stancofski, Chloe Adrienna Talana, I-Ting Teng, Shuishu Wang, Kizzmekia S. Corbett, Yaroslav Tsybovsky, John R. Mascola, Peter D. Kwong

**Affiliations:** Vaccine Research Center, National Institute of Allergy and Infectious Diseases, National Institutes of Health, Bethesda, MD 20892, USA; Electron Microscopy Laboratory, Cancer Research Technology Program, Leidos Biomedical Research, Inc., Frederick National Laboratory for Cancer Research, Frederick, MD 21702, USA

**Keywords:** beta-coronavirus, broadly neutralizing antibody, COVID-19, crystal structure, SARS-CoV-2, S2-directed antibody, vaccine design

## Abstract

Immunization with SARS-CoV-2 spike elicits diverse antibodies, but can any of these neutralize broadly? Here, we report the isolation and characterization of antibody WS6, from a mouse immunized with mRNA encoding the SARS-CoV-2 spike. WS6 bound diverse beta-coronavirus spikes and neutralized SARS-CoV-2 variants, SARS-CoV, and related sarbecoviruses. Epitope mapping revealed WS6 to target a region in the S2 subunit, which was conserved among SARS-CoV-2, MERS-CoV, and hCoV-OC43. The crystal structure at 2-Å resolution of WS6 with its S2 epitope revealed recognition to center on a conserved helix, which was occluded in both prefusion and post-fusion spike conformations. Structural and neutralization analyses indicated WS6 to neutralize by inhibiting fusion, post-viral attachment. Comparison of WS6 to other antibodies recently identified from convalescent donors or mice immunized with diverse spikes indicated a stem-helical supersite – centered on hydrophobic residues Phe1148, Leu1152, Tyr1155, and Phe1156 – to be a promising target for vaccine design.

**Highlights:**

- SARS-CoV-2 spike mRNA-immunized mouse elicited an antibody, WS6, that cross reacts with spikes of diverse human and bat beta-coronaviruses
- WS6 neutralizes SARS-CoV-2 variants, SARS-CoV, and related viruses
- Crystal structure at 2-Å resolution of WS6 in complex with a conserved S2 peptide reveals recognition of a helical epitope
- WS6 neutralizes by inhibition of fusion, post-viral attachment
- WS6 recognizes a supersite of vulnerability also recognized by other recently identified antibodies
- Helical supersite of vulnerability comprises a hydrophobic cluster spanning three helical turns, with acid residues framing the center turn
- Genetic and structural analysis indicate supersite recognition to be compatible with diverse antibody ontogenies

## Introduction

The COVID-19 pandemic, resulting from the zoonotic infection of severe acute respiratory syndrome coronavirus 2 (SARS-CoV-2), continues to rage on, fueled by continuously evolving variants, which are making current licensed vaccines less effective (Araf et al., 2022; Doria-Rose et al., 2021; Edara et al., 2021; Garcia-Beltran et al., 2022; Liu et al., 2021). Vaccines capable of neutralizing all SARS-CoV-2 variants for the foreseeable future are of high interest. Antibodies with broad neutralizing capacity are also of interest: if ultrapotent, they might be useful as therapeutic antibodies; but even if only moderate potency, their epitopes are useful as vaccine templates (Kong et al., 2016).

Virtually all neutralizing antibodies are directed against the trimeric ectodomain of the spike glycoprotein, which comprises two subunits S1 and S2. Neutralizing antibodies isolated from COVID-19 convalescent donors or from vaccinees after spike immunization are directed primarily against the N-terminal domain (NTD) or receptor-binding domain (RBD) on the S1 subunit of the trimeric viral surface spike glycoprotein (Spike) (Barnes et al., 2020; Brouwer et al., 2020; Cao et al., 2020; Cerutti et al., 2021; Ju et al., 2020; Liu et al., 2020; McCallum et al., 2021; Robbiani et al., 2020; Rogers et al., 2020; Seydoux et al., 2020; Suryadevara et al., 2021; Zost et al., 2020). Evolving SARS-CoV-2 variants, such as Delta and Omicron, evade these antibodies by mutations that reduce or knockout antibody binding, but maintain or even enhance infectivity (Garcia-Beltran et al., 2022; Liu et al., 2021; Sievers et al., 2022; Syed et al., 2022). Antibodies against most other regions on the spike are generally poorly neutralizing to non-neutralizing; several antibodies, however, such as antibody S2P6 (Pinto et al., 2021) have been reported to neutralize diverse strains of beta coronaviruses through recognition of a stem-helix supersite of vulnerability in the S2 subunit (Hsieh et al., 2021; Li et al., 2022; Sauer et al., 2021; Zhou et al., 2021).

To investigate the breadth of neutralizing antibodies obtained from mice vaccinated by mRNA encoding the SARS-CoV-2 spike, we assessed monoclonal antibodies for the location of their epitopes, the breadth of their binding to diverse spikes, and their neutralization capacities. We found one, antibody WS6, with broad binding capacity and moderate neutralization potency, and we determined its crystal structure in complex with its epitope, the step in the entry pathway where it neutralized, and how its recognition compared with other recently identified antibodies with overlapping epitopes. The results reveal a highly promising vaccine target in the S2 subunit – comprising a hydrophobic cluster spanning three helical turns, with acidic residues framing its center turn – and add WS6 to the panel of antibodies by which to guide its vaccine development.

## Results

### Identification and characterization of SARS-CoV-2 spike-specific antibodies from immunized mice

To obtain antibodies specific for SARS-CoV-2 spike glycoprotein, we immunized mice with mRNA coding for SARS-CoV-2 spike (**Figure 1A**). To generate hybridomas, we boosted with soluble spike protein and after three days generated hybridomas by fusing splenocyte B cells with Sp2/0 cells from the mouse with the highest plasma neutralization titers to SARS-CoV-2. Eleven monoclonal antibodies, named WS1 to WS11, bound SARS-CoV-2 S-dTM by ELISA (**Figure 1B**). Nine of these bound the S1 subunit, either S1-short1 (spike residues 1-670) or S1R (residues 1-537). Six of them, WS1, WS2, WS3, WS7, WS8, and WS10, bound NTD; and three of them, WS4, WS9, and WS11, bound RBD. Antibodies WS5 and WS6, however, did not bind NTD, RBD, or S1, and their binding epitopes were presumably on the S2 subunit of the spike.

**Figure 1.**
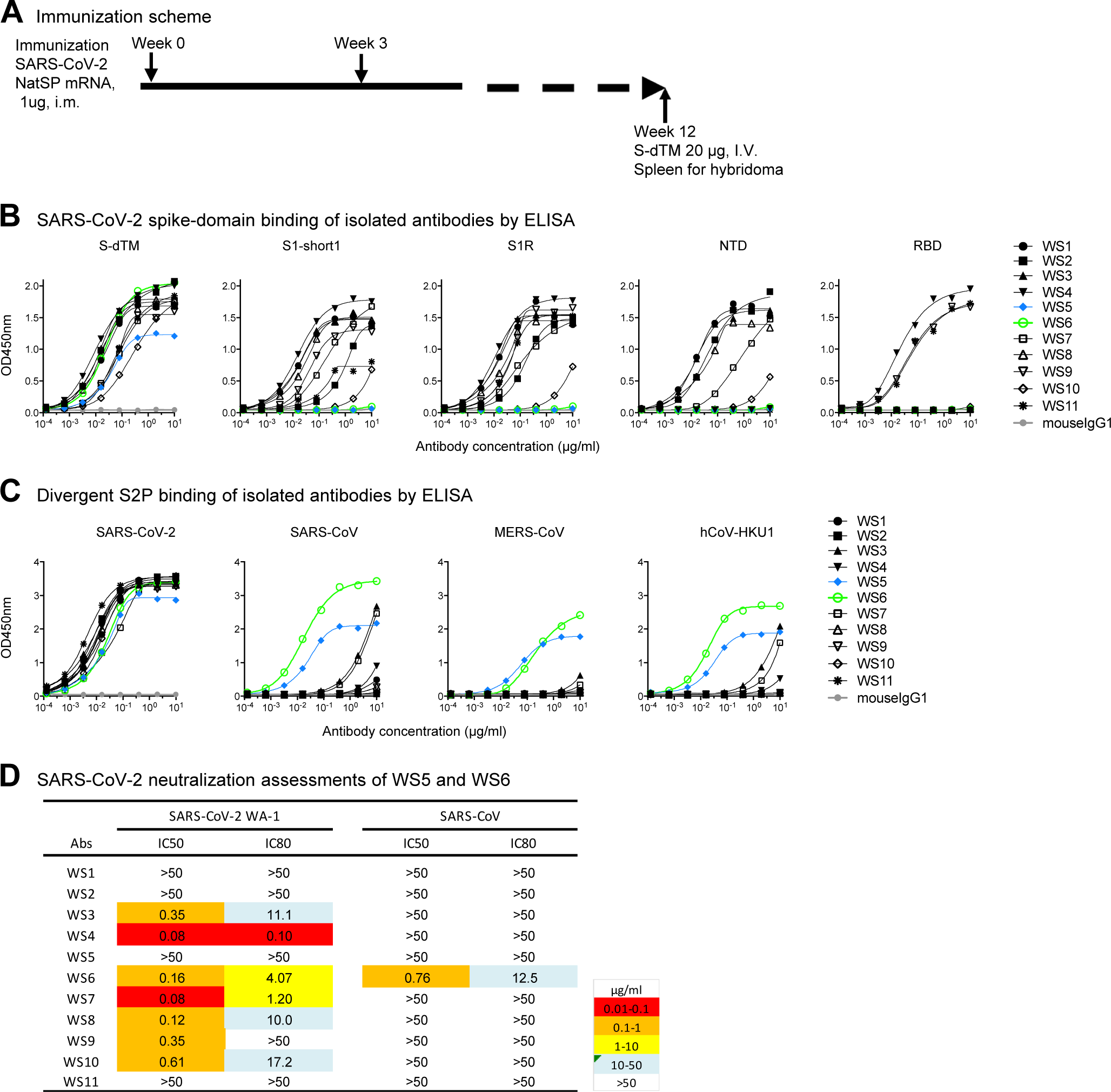
Spike mRNA immunized mice elicit antibodies against diverse regions of spike, several of which bound diverse beta-coronavirus spikes and one of which, WS6, neutralized. **(A)** Immunization scheme. NatSP is the full transmembrane-containing native sequence of spike WA-1 strain; S-dTM is the soluble spike protein residues 1-1206 of wildtype WA-1 strain. **(B)** Binding of isolated antibodies by ELISA to prefusion-stabilized spike (S2P) and its subdomains. Purified monoclonal antibodies from hybridoma supernatants were analyzed for binding to SARS-CoV-2 S-dTM, S1 (S1-short and S1R), RBD and NTD by ELISA. **(C)** Binding of isolated antibodies assess by ELISA to diverse beta-coronavirus prefusion-stabilized spikes (S2P). **(D)** Neutralization assessment of hybridoma antibodies against SARSCoV-2 WA-1 and SARS-CoV pseudovirus on 293T-ACE2 cells. See also Figure S1.

To provide insight into the breadth of binding, we assessed recognition of WS1-11 on a panel of prefusion-stabilized diverse beta-coronavirus spikes (S2Ps) (Wrapp et al., 2020). Detectable binding was observed against SARS-CoV-2, SARS-CoV, MERS-CoV and hCoV-HKU1 for five antibodies (**Figure 1C**). For antibodies WS3, WS4, and WS7, binding was more than 1,000-times weaker against the divergent strain versus that of the immunogen. However, for antibodies WS5 and WS6, binding was only about 10-fold reduced versus SARS-CoV-2. Neutralization assessments revealed that WS5 neutralized neither SARS-CoV-2 nor SARS-CoV, whereas WS6 could neutralize both (**Figure 1D**). WS6 was further studied for its epitope and neutralization activities.

### WS6 recognizes diverse beta-coronaviruses spikes and neutralizes SARS-CoV-2 and variants, SARS-CoV, and related viruses from bat, pangolin, and other animals

To assess fully the broad reactivity of WS6, we performed ELISAs against S2Ps from an even more divergent panel of coronaviruses, including RaTG13, WIV1, SHC014, hCoV-OC43, and hCoV-229E (**Figure 2A)**. Remarkably, WS6 was able to bind all beta-coronaviruses tested, though not to the alpha-coronavirus hCoV-229E **(Figure S1**). The apparent binding affinities of WS6 to S2Ps, measured by biolayer interferometry (BLI), showed nanomolar or lower dissociation constants (K_D_s), with slow dissociation such that k_off_ values could not be determined accurately in many cases (**Figure 2B**). To determine if WS6 could bind spike proteins on cell surface, we expressed spikes of SARS-CoV-2, SARS-CoV, MERS-CoV, hCoV-HKU1, and hCoV-OC43 on the surface of Expi-293 cells and analyzed WS6 binding by flow cytometry. We found that WS6 bound well to the cell surface expressing the beta-coronavirus spikes (**Figures 2C and S2**).

**Figure 2.**
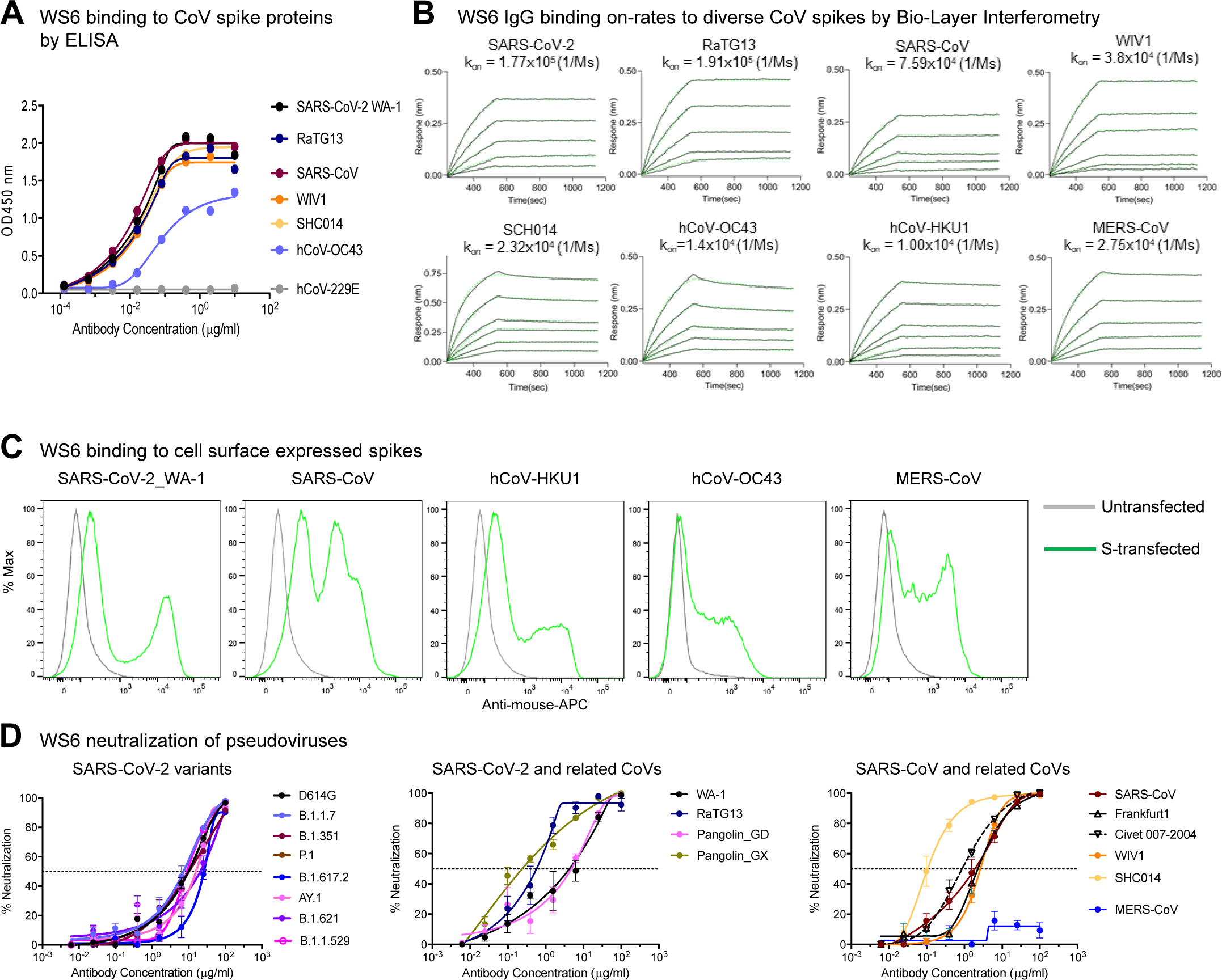
Antibody WS6 binds and neutralizes diverse beta-coronaviruses. **(A**) ELISA binding analysis of WS6 to prefusion-stabilized soluble spikes of various coronaviruses. **(B)** BLI binding curves of WS6 with various beta-coronaviruses. **(C)** WS6 binds cell surface-expressed spike proteins from SARS-CoV-2, SARS-CoV, hCoV-HKU1, hCoV-OC43 and MERS-CoV. **(D)** WS6 neutralizes SARS-CoV-2 variants, SARS-CoV and related animal coronaviruses. Neutralization activity was measured using spike-pseudotyped lentivirus on 293 flpin-TMPRSS2-ACE2 cells. Assays were performed in triplicate on 293 flpin-TMPRSS2-ACE2 cells, and representative neutralization curves from 2-3 technical replicates are shown. (Left) WS6 neutralizes SARS-CoV-2 variants, D614G, B.1.1.7 (Alpha), B.1.351 (Beta), P.1 (Gamma), B.1.617.2 (Delta), AY.1 (Delta+), B.1.621 (Mu) and B.1.1.529 (Omicron). (Middle) WS6 neutralizes SARS-CoV-2 and related coronaviruses, SARS-CoV-2 WA-1, RaTG13, Pangolin_GD and Pangolin_GX. (Right) WS6 neutralizes SARS-CoV and related coronaviruses, SARS-CoV, Frankfurt1, Civet 007-2004, WIV1 and SHC014, but not MERS-CoV. See also MERS-CoV. See also Figure S1, S2, S3 and Table S1. Figure S1, S2, S3 and Table S1.

To determine if the broad recognition of WS6 for spikes translated into broad neutralization of beta-coronaviruses, we assessed its neutralization activities by pseudovirus neutralization assays (Naldini et al., 1996; Wang et al., 2021). WS6 neutralized all tested variants of SARS-CoV-2 including Omicron (B.1.1.529), with IC_50_ 2.46-26.52 μg/ml (**Figures 2D and S3; Table S1**). WS6 could also neutralize beta-coronaviruses related to SARS-CoV-2 (such as RaTG13, Pangolin_GD, and Pangolin_GX), SARS-CoV, and related viruses Frankfurt1, Civet007-2004, WIV1, and SHC014 with IC_50_ 0.11-4.91 μg/ml (**Figures 2D and S3; Table S1**). WS6 also neutralized related beta-coronaviruses, such as RaTG13, Pangolin-GX, Civet007-2004, and SHC014, with sub micromolar IC_50_, better than against SARS-CoV-2 despite being elicited by immunizations with SARS-CoV-2 spike. The only tested beta-coronavirus strain WS6 failed to neutralize was MERS-CoV, consistent with its lower ELISA binding to this strain (**Table S1**).

### WS6 epitope mapping

We attempted to map the epitope of WS6 by visualizing its recognition of the spike ectodomain by negative strain-electron microscopy (EM). 2D-classification of antigen-binding fragment (Fab) of WS6 in complex with the S2P spike (WA-1 strain) showed generally unbound spikes, with only 4% of the images yielding a trimer with Fabs binding in the membrane-proximal S2 stem (**Figure 3A**). We also analyzed Fab WS6 in complex with spike S2 subunit; 2D-classification indicated WS6 to bind S2, though most of S2 appeared to be disordered (**Figure 3B**).

**Figure 3.**
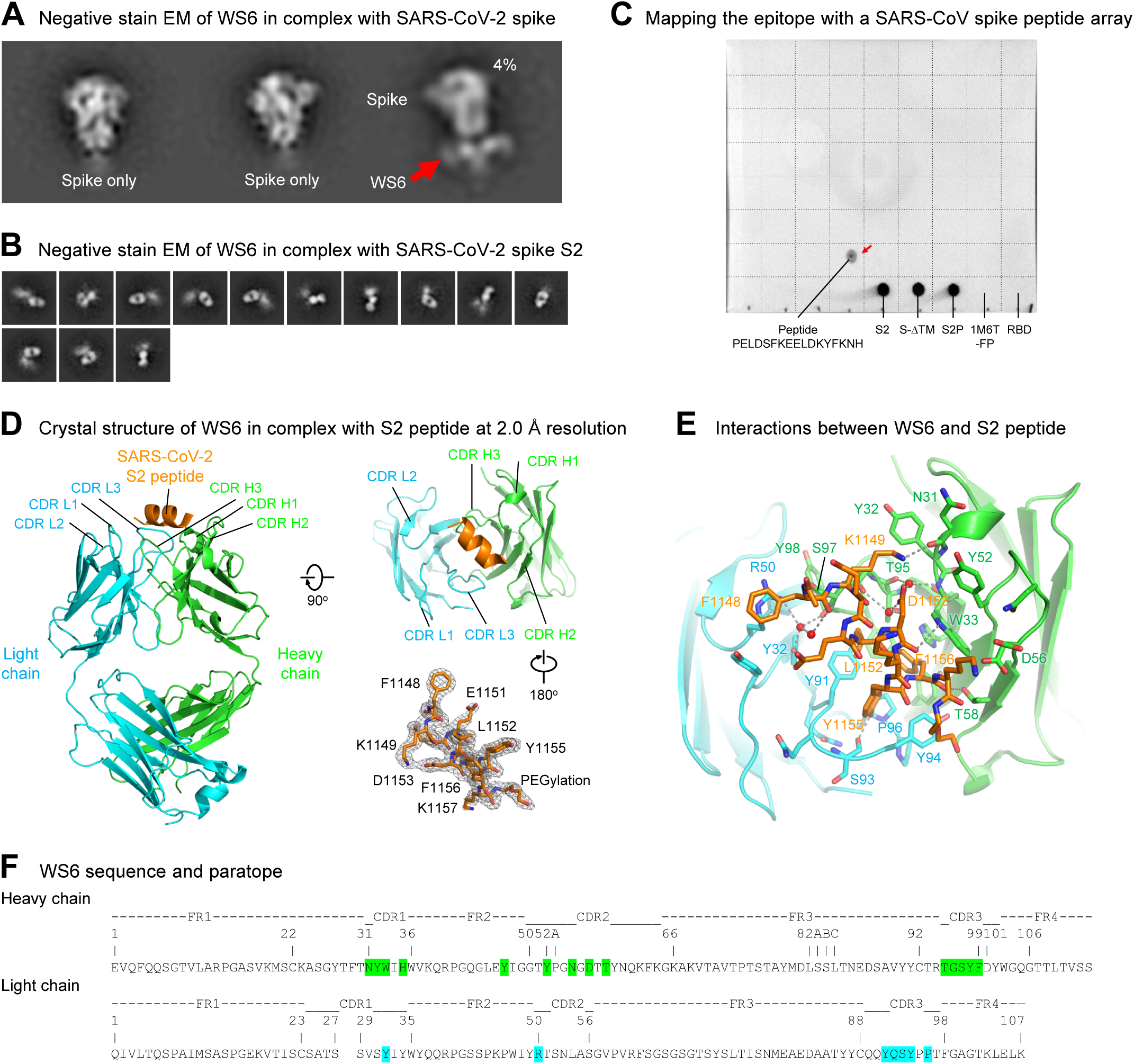
Epitope mapping and crystal structure of antibody WS6 in complex with S2 peptide. **(A)** Negative stain EM of SARS-CoV-2 spike in complex with WS6. Only a small fraction was observed with WS6 bound at the stem region. **(B)** Negative stain EM of SARS-CoV-2 spike S2 in complex with WS6. **(C)** Epitope mapping by dot-blot assay with a SARS-CoV spike peptide array (BEI, NR-52418). WS6 bound to peptide #154 P1125ELDSFKEELDKYFKNH1141 (corresponding to SARS-CoV-2 residues 1143-1159) (red arrow), SARS-CoV-2 spike and its S2 domain. A dash line grid was added on top of the blot to aid display. **(D)** Crystal structure of WS6 in complex with SARS-CoV-2 S2 peptide at 2.0 Å resolution. The antibody and SARS-CoV-2 peptide are shown in cartoon representation. WS6 heavy chain, light chain and S2 peptide are colored green, cyan and orange, respectively. Two 90o-flipped views were shown (left, top right). Electron density map of the S2 peptide was shown with the antibody interacting residues facing the reader (right, bottom) in a 180o-flipped view from panel above. **(E)** Detailed interactions between WS6 and S2 peptide. The S2 peptide and paratope residues in WS6 were shown in sticks representation with other regions of WS6 shown in cartoon representation. Hydrogen bonds were indicated with gray dashed lines. Waters that mediating hydrogen bonds were shown as red spheres. **(F)** Sequence and paratope of WS6. WS6 residues were numbered according to Kabat nomenclature. Heavy and light chain paratope residues were highlighted in green and cyan, respectively. See also Table S3.

To further map the epitope, we performed peptide array-based epitope mapping and identified a 17-residue peptide, PELDSFKEELDKYFKNH (SARS-CoV-2 spike residues 1143-1159) to bind WS6 (**Figure 3C**), suggesting WS6 epitope to be at the S2 stem-helix region of the spike near the viral membrane, consistent with our negative stain-EM observation. This peptide is conserved among SARS-CoV, SARS-CoV-2, and RaTG13, and is mostly conserved among diverse beta-coronaviruses (**Figure S4A**).

### Crystal structure of WS6 in complex with a conserved S2 peptide

To elucidate the mechanism for the broad reactivity of WS6, we determined a crystal structure at 2 Å of Fab WS6 in complex with peptide Ace-F_1148_KEELDKYFK_1157_-PEG12-Lys-Biotin (residue numbers were based on the SARS-CoV-2 spike sequence) (**Figure 3D and Table S2**). The peptide contained a 10-residue segment conserved among beta-coronaviruses (**Figure S4A**) and was acetylated at N terminus and biotin-pegylated at C terminus. Well-defined electron density was observed for the entire WS6 Fab and for the entire peptide, including the acetyl group at the N terminus and part of the polyethylene glycol at the C terminus (**Figure 3D**). Peptide binding interactions involved both heavy and light chains, with heavy chain contributing ∼360 Å^2^ buried surface area (BSA) and light chain contributing ∼230 Å^2^ (**Table S3**). The BSA of the peptide was slightly larger at ∼675 Å^2^, not counting the BSA of the visible PEG fragment (∼100 Å^2^ total). This binding interface is smaller than a typical antibody-binding epitope, indicating the full WS6 epitope to likely involve additional residues.

All complementarity-determining regions (CDRs) of both heavy and light chains were involved in binding, creating a groove that cradled the epitope (**Figure 3D**). In the WS6-bound crystal structure, the S2 epitope formed a three-turn α helix; examination of its binding mode revealed binding for a longer helix with extensions on both termini without any major clashes. Most of the residues of the peptide involved in binding, except for Glu1150 and Lys1154, which were on the side of the helix facing WS6 (**Figure 3E**). Aromatic or hydrophobic residues, Phe1148, Leu1152, Tyr1155, and Phe1156, were on the side of the helix facing WS6. Phe1148 had aromatic interactions with Arg50_L_ and Tyr91_L_ of WS6 light chain and hydrophobic interactions with Ser97_H_ of heavy chain. Tyr1155 side chain stuck into a cavity form by CDR L3 from Tyr91_L_ to Pro96_L_ and had aromatic or hydrophobic interactions with side chains of Tyr91_L_, Tyr94_L_, and Pro96_L_ and the peptide backbone. Leu1152 and Phe1156 side chains bound in a pocket between heavy and light chains and interacted with side chains of Tyr91_L_, Pro96_L_, Phe99_H_, Thr95_H_, Trp33_H_, His35_H_, and Tyr47_H_ (**Figure 3E and 3F and Table S3**).

### WS6 neutralizes by inhibition of fusion steps post viral attachment

To elucidate the mechanism by which WS6 neutralize beta-coronaviruses, we incubated BHK21-ACE2 cells with SARS-CoV-2 spike pseudotyped lentivirus on ice to allow virus to attach to ACE2. After thorough washing, cells were incubated with WS6 or WS4 on ice for one hour and at 37 L for 72 hours. WS4, an RBD-directed neutralizing antibody, could neutralize not more than 40% of the virus, whereas WS6 had no problem neutralizing the ACE2 pre-attached virus (**Figure 4A**). We performed a similar experiment using 293 flpin-TMPRSS2-ACE2 cells for WS6 and S2P6 (Pinto et al., 2021), which also binds to the S2-stem helix, and found that both antibodies could neutralize ACE2 pre-attached virus. These results suggest that WS6 is likely to neutralize SARS-CoV-2 by engaging in steps post-viral attachment to ACE2.

**Figure 4.**
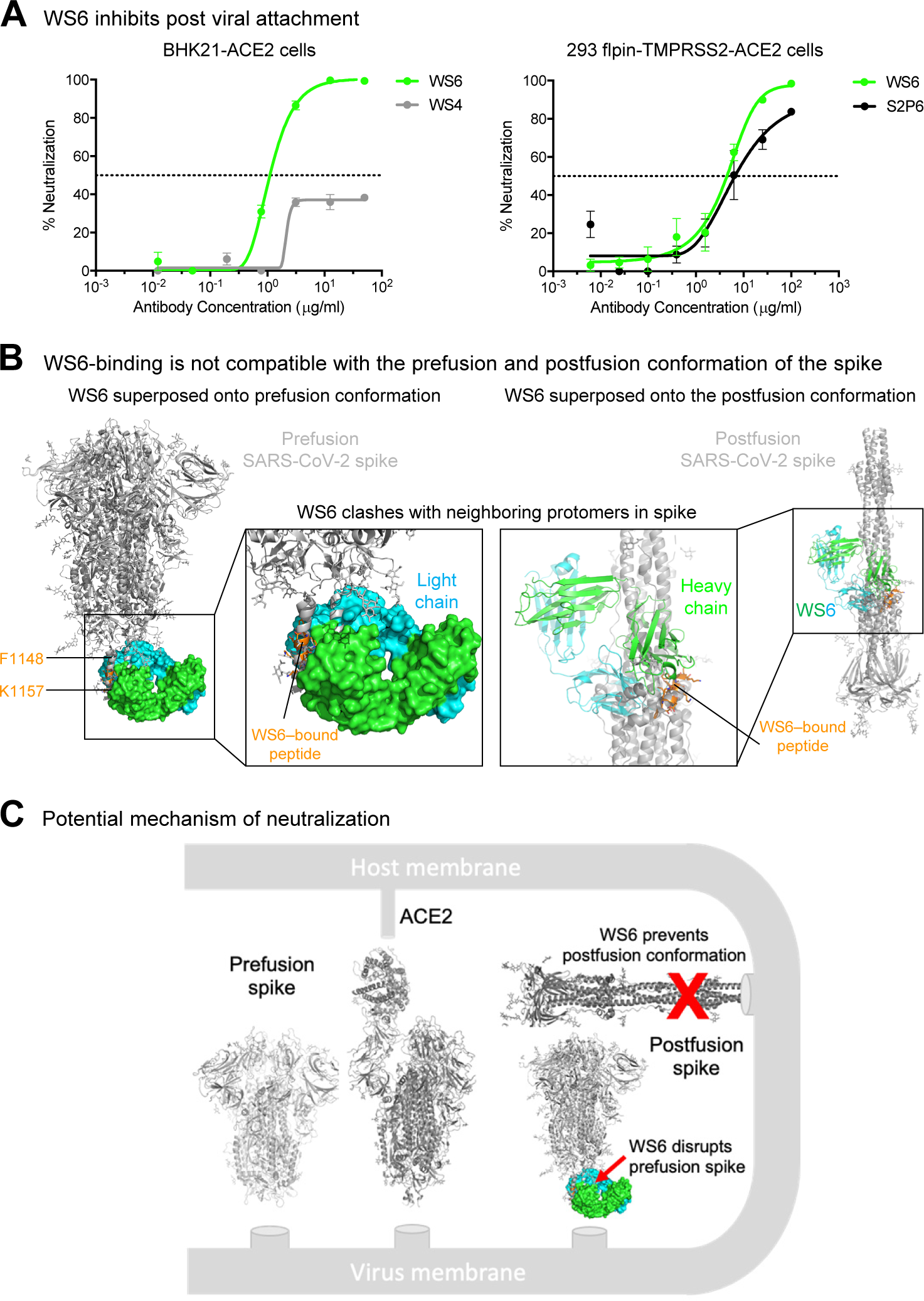
WS6 neutralizes SARS-CoV-2 by inhibition of post viral attachment. **(A)** WS6 and S2P6 neutralization of SARS-CoV-2 by inhibition of post viral attachment. Neutralization experiments were performed with two BHK21-ACE2 (top) and 293 flpin-TMPRSS2-ACE2 (bottom) cells. WS4, a RBD antibody, was used as a negative control. Representative neutralization curves from two technical replicates of experiments are shown. **(B)** Binding mode of WS6 is not compatible with the prefusion and postfusion conformation of the spike. The WS6-S2 peptide complex was superposed onto the pre- and post-fusion spike trimer by aligning the WS6-bound S2 peptide with the corresponding part in one of the protomers. S2-bound WS6 clashed with neighboring protomers in either prefusion or postfusion conformation, suggesting binding disrupts prefusion conformation and prevents formation of postfusion. **(C)** Mechanism of neutralization by WS6. Structure of WS6 in complex with S2 peptide indicated that WS6 disrupts prefusion spike and prevents formation of the postfusion conformation.

As described above in the structural analysis, WS6 recognized the hydrophobic face of the stem helix with residues Phe1148, Leu1152, Tyr1155, and Phe1156 binding in the center of the paratope groove. In the prefusion structure of SARS-CoV-2 spike (PDB: 6xr8) (Cai et al., 2020; Ke et al., 2020), these hydrophobic side chains pack in the coiled-coil interface of the 3-helical bundle. Modeling of WS6 binding to the prefusion SARS-CoV-2 spike, based on the WS6-peptide complex, revealed substantial clashes (**Figure 4B, left panel**), indicating that WS6 binding would require unpacking or disassembly of the helical bundle. Similarly, the postfusion spike structure (PDB: 6xra) (Cai et al., 2020) was incompatible with binding of WS6 to the hydrophobic side of the peptide helix as revealed by the crystal structure, because these hydrophobic residues are involved in the packing of the small helix with the central coiled coil in the postfusion structure (**Figure 4B, right panel**). Because the spike binding to the receptor and transitioning from the prefusion conformation to postfusion conformation drive the membrane fusion process, WS6 maybe even more apt to bind during this conformational transition and inhibit the fusion process (**Figure 4C**).

### A helical stem supersite of vulnerability

Several broad beta-coronavirus-neutralizing antibodies have been identified recently that target the S2 stem-helix region, including the afore mentioned S2P6 (Pinto et al., 2021) as well as antibodies B6 (Sauer et al., 2021), IgG22 (Hsieh et al., 2021), CCP40.8 (Zhou et al., 2021), and CV3-25 (Li et al., 2022). Superposition of their S2 helical epitopes indicated these antibodies to bind to the stem helix with varying orientations (**Figure 5A**) and to target different sets of residues ranging from residue position 1142 to 1164 on S2 (**Figures 5B and S5**). Epitope analysis indicated a common subset of residues, namely Phe1148, Lys1149, Glu1153, Leu1152, Asp1153, Tyr1155, and Phe1156, to interact with four of the six antibodies (**Figure 5B, left**). These residues, which spanned three helical turns on S2 with a central hydrophobic cluster sandwiched by hydrophilic residues Lys1149, Glu1151, and Asp1153, formed a supersite of vulnerability for antibody recognition (**Figure 5B, middle and right**). Sequence analysis indicated this S2 supersite to be highly conserved among beta-coronaviruses (**Figure 5C**), providing the basis for the broad neutralization by WS6 and other antibodies targeting this site. Overall, these S2 stem-helix antibodies are likely to share a similar mechanism of neutralization. Before or even after spike binds to ACE2, the stem helix appears to adopt a conformation or conformations that expose this supersite, allowing for the stem-helix antibodies to bind and prevent spike conformations needed for fusion, thereby stalling the entry process. Importantly, these S2-helix directed broad neutralizers have diverse origin genes, except for B6 and IgG22, which utilize the same VH gene and appears to be of the same antibody class (Figure S4B). In addition, the antibodies have only low-to-moderate somatic hypermutation, suggesting diverse ontogenies are possible with little barrier to their development **(Figure S4)**.

**Figure 5.**
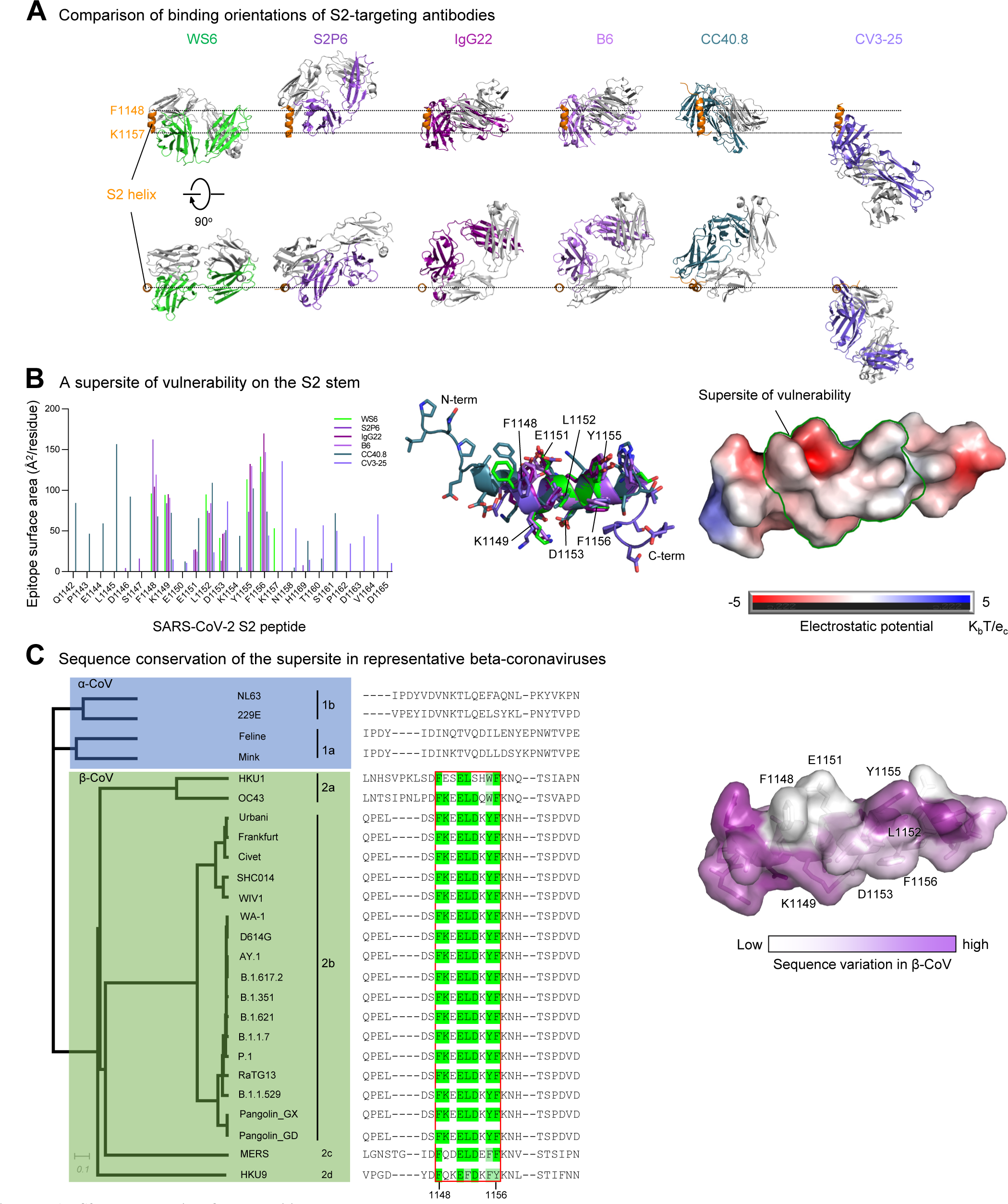
An S2-stem supersite of vulnerability. **(A)** Comparison of modes of recognition of WS6 with other antibodies targeting the S2 stem helix. WS6 binding to the S2 helix was shown in the same orientation as in Fig. 4B. S2P6, IgG22, B6, CC40.8 and CV3-25 complexes were aligned with the WS6 complex over the S2 peptide helix. The S2 helix and light chains of antibodies were colored orange and gray, respectively. Heavy chains of antibodies are colored differently to distinguish. CV3-25 assumes a distinct mode of recognition comparing to others. **(B)** Epitopes of antibodies targeting the S2 region define a supersite of vulnerability. Antibody-binding surface areas of each residue in the S2 peptide were plotted along the linear sequence of SARS-CoV-2 S2 peptide (left). Antibody-bound peptides were shown colored by recognizing antibodies. The center region between residues 1148 and 1156 were recognized by most of the antibodies (middle), hydrophobic residues F1148, L1152, Y1155 and F1156 were positioned in the middle of the supersite with K1149, E1151 and D1153 providing hydrophilic interactions are the peripheral (right). The boundary of the supersite, which was defined as residues interacting with 5 of the 6 antibodies analyzed in this study, was highlighted by a green line. **(C)** Sequence conservation of the SARS-CoV-2 supersite inβ-coronaviruses. Sequence dendrogram was calculated using full spike protein sequences (far left). Sequence alignment shown residues 1142 – 1165, recognized by supersite antibodies in panel B (left middle). The surface of S2 peptide was colored by sequence variation score with corresponding residues shown in sticks representation (right). See also Figures S4 and S5.

## Discussion

Zoonotic infections from beta-coronaviruses have caused multiple pandemics and endemics in recent years, including SARS (Drosten et al., 2003; Ksiazek et al., 2003), Middle East respiratory syndrome (MERS) (Zaki et al., 2012), and the ongoing COVID-19 (Zhou et al., 2020a; Zhu et al., 2020). It seems likely that additional beta-coronavirus zoonotic pandemics will occur in the future. Broadly neutralizing antibodies or broad vaccines against a wide spectrum of beta-coronaviruses may be effective at preventing or ameliorating such pandemics – and could even be of use against the current COVID-19, if variants do succeed in escape control by current vaccines. In this study, we isolated an S2-directed antibody, named WS6, from a mouse immunized with mRNA-encoded SARS-CoV-2 spike. WS6 could bind and neutralize all variants of SARS-CoV-2, including Delta and Omicron; it could also neutralize SARS-CoV, as well as bat, civet, and pangolin beta-coronaviruses related to SARS-CoV and SARS-CoV-2. Crystal structure of WS6 in complex with S2 stem-helix epitope peptide revealed the structural basis for broad recognition and, together with neutralization and binding analyses, suggest a potential mechanism by which WS6 neutralize broadly beta-coronaviruses.

Several other groups have recently reported S2-directed antibodies including antibodies S2P6 (Pinto et al., 2021), CC40.8 (Zhou et al., 2021) and CV3-25 (Li et al., 2022) from SARS-CoV-2-infected convalescent donors and B6 (Sauer et al., 2021) and IgG22 (Hsieh et al., 2021) from vaccinated mice. We measured directly the neutralization of both WS6 and S2P6 to diverse SARS-CoV-2 variants and to diverse coronaviruses generally (**Figure S3**). In light of its high breadth, we were surprised to find S2P6 was unable to neutralize the Omicron variant of SARS-CoV-2, although WS6 did neutralize Omicron. WS6 did, however, neutralize all tested strains more potently than S2P6, except for MERS-CoV, against which WS6 was non-neutralizing. In terms of the murine antibodies, the recognition of antibodies B6 and IgG22 was very similar, both utilizing the same heavy chain origin gene (VH1-19) and both generated after immunization with spikes from SARS-CoV-2 and MERS-CoV (Hsieh et al., 2021; Pinto et al., 2021). By contrast, WS6 did not use the same V-gene and was generated only by immunization with the SARS-CoV-2 spike – thereby revealing a new mode of murine S2-helix recognition and showing MERS-CoV spike immunization was no required to elicit these antibodies.

It will be interesting to determine whether S2-helix peptide-based immunizations can focus the immune response, enabling the elicitation of broadly neutralizing serological responses, as has been done with HIV-1 fusion peptide (Kong et al., 2019; Xu et al., 2018). Nanoparticles displaying the S2-helix may also be helpful in focusing the immune response, as has been done with the influenza stem cite of vulnerability (Boyoglu-Barnum et al., 2021; Kanekiyo et al., 2013; Yassine et al., 2015). Whether S2-helix focused immunization will enable sufficiently broad and potent responses to enable protection from future SARS-CoV-2 variants or to future beta-coronavirus zoonotic crossovers remains to be seen. It seems likely that such immune focusing can be carried out in conjunction with spike-based immunizations, perhaps using currently licensed vaccines from Moderna, Pfizer and Johnson and Johnson, all of which incorporate the S2-helical region.

## Acknowledgements

We thank R. Andrabi for providing CC40.8 coordinates, B. Graham for suggesting use of flpin-TMPRSS2-ACE2 cells, T. Stephens for assistance with negative-stain EM, J. Stuckey for assistance with figures, and members of the Virology Laboratory, Vaccine Research Center, for discussions and comments on the manuscript. Support for this work was provided by the Intramural Research Program of the Vaccine Research Center, National Institute of Allergy and Infectious Diseases, National Institutes of Health and by federal funds from the Frederick National Laboratory for Cancer Research under Contract HHSN261200800001E. Use of sector 22 (Southeast Region Collaborative Access team) at the Advanced Photon Source was supported by the US Department of Energy, Basic Energy Sciences, Office of Science, under contract number W-31-109-Eng-38.

## Author Contributions

W.S. isolated antibodies WS1-WS11 and performed ELISA binding assays; L.W. headed neutralization studies, prepared spike plasmids and performed cell surface binding assays; T.Z. determined and analyzed the crystal structure of WS6-peptide complex, and wrote manuscript; M.S. performed Octet binding assays, provided spike proteins for ELISA, and assisted with obtaining peptide for crystallization; E.S.Y., Y.Z., and M.Chen performed WS6 neutralization of SARS related viruses; X.C. provided mouse NGS sequence data; M.Choe provided S2P6 antibody; A.C. provided 293 flpin-TMPRSS2-ACE2 cells; K.L. assisted with preparation of pseudoviruses; A.S.O provided SARS-CoV-2 S2P-His for Octet measurements; A.P. and C.A.T performed cell surface staining; R.R. assisted with obtaining Peptide for crystallization; C.-H.S. prepared phylogenetic tree; E.-S.S. crystallized the protein complex; I.-T.T. performed transfection for Beta-CoV and S2 antibody productions; S.W. wrote manuscript and performed PISA analysis of antibody-peptide interface; K.S.C carried out mouse immunization experiment; Y.T. performed negative-stain EM analysis; J.R.M. supervised antibody isolation, neutralization assessments, and cell surface binding assays; P.D.K. oversaw the project and - with S.W and T.Z - wrote the manuscript, with all authors providing comments or revisions.

## METHODS

### RESOURCE AVAILABILITY

#### Materials Availability

Plasmids generated in this study are available upon request.

#### Data and Code Availability

Crystal diffraction data and structure coordinates of WS6 in complex have been deposited with the Protein Data Bank, PDB: 7TCQ, DOI: 10.2210/pdb7tcq/pdb.

### EXPERIMENTAL MODEL AND SUBJECT DETAILS

#### Mouse Studies

Animal experiments were carried out in compliance with all pertinent US National Institutes of Health regulations and approval from the Animal Care and Use Committee (ACUC) of the Vaccine Research Center. BALB/cJ mice (Jackson Laboratory), 6- to 8-week-old female, were used.

#### Cell Lines

FreeStyle 293-F (cat# R79007) and Expi293F cells (cat# A14528; RRID: CVCL_D615) were purchased from ThermoFisher Scientific Inc. FreeStyle 293-F cells were maintained in FreeStyle 293 Expression Medium, while Expi293F cells were maintained in Expi Expression Medium. The above cell lines were used directly from the commercial sources and cultured according to manufacturer suggestions.

### METHOD DETAILS

#### Mouse immunization, hybridoma generation and mAb isolation

Mice were immunized with mRNA coding for SARS-CoV-2 spike of the wildtype WA-1 strain at weeks 0 and 3 intramuscularly, utilizing methods described previously (Corbett et al., 2020). One mouse with the best neutralizing antibody titer against SARS-CoV-2 spike was boosted intravenously with 20□μg of SARS-CoV-2 S-dTM at week 12. Three days later, splenocytes were harvested and fused with Sp2/0 myeloma cells (ATCC) using polyethylene glycol (PEG) 1450 (50% (w/v), Sigma-Aldrich) according to the standard methods. Cells were cultured and screened in RPMI complete medium that contained 20% FCS and 1 × 100□μM hypoxanthine, 0.4□μM aminopterin and 16□μM thymidine (Sigma-Aldrich). Supernatants from resulting hybridomas were screened for binding, using ELISA, to SARS-CoV-2-S1, NTD, RBD or S-dTM as well as for neutralizing activity. Subclones were generated by limiting dilution. After three rounds of screening and subcloning, stable antibody-producing clones were isolated and adapted to hybridoma-serum-free medium (Life Technologies Corp., Grand Island, NY, USA). Supernatants were collected from selected hybridoma clones and purified through a protein G-sepharose column (GE Healthcare). mAbs were isotyped with Pierce rapid isotyping kit (Cat#26178). mAb Fabs were generated using Pierce Fab kit (Cat#44985) following manufacturer’ s instructions. mAb and Fab purity was confirmed by SDS-PAGE. Selected hybridoma were sent to GenScript (Piscataway, NJ 08854, USA) for hybridoma heavy and light chain variable sequences and IMGT analysis.

#### ELISA and dot-blot

ELISA plates were coated with the SARS-CoV-2 proteins, S-dTM, S1-short (residues 1-670), S1R (residues 1-537), RBD, and NTD, at 1□μg/ml in PBS at 4□°C overnight. After standard washes and blocks, plates were incubated with 50ul/each well, serial dilutions of mAbs for one hour at room temperature. Anti-mouse whole IgG horseradish peroxidase conjugates (Jackson Laboratory) were used as secondary antibodies, and 3,5,3′5′-tetramethylbenzidine (TMB) (KPL, Gaithersburg, MD) was used as the substrate.

To identify WS6 epitope, a dot-blot was performed. SARS-CoV spike peptide array (NR-52418), was purchased from BEI resource. Seventy-eight peptides (#91-169) from the peptide array which covers SARS-CoV spike S2 region, were dotted on a nitrocellulose membrane respectively, 20ng/2ul/each dot. SARS-CoV-2 RBD, S2, S-dTM and S2P proteins were also dotted on the membrane as controls. Block non-specific sites by soaking in 5% dry milk in PBS-Tween 20. The membrane was washed three times with PBS-T buffer, then incubated with 2ug/ml of mAb, WS6 in blocking buffer for one hour at room temperature. After wash three times with PBS-T buffer, the membrane was incubated half hour with anti-mouse IgG horseradish peroxidase conjugates (Jackson Laboratory) as secondary antibody. Then membrane was developed with ECL medium.

#### Expression and purification of beta-coronavirus spike trimer proteins

Diverse beta-coronavirus spike soluble proteins were stabilized in prefusion conformation by double-proline mutations corresponding to K986P and V987P in SARS CoV-2 spike protein, along with a T4-phage fibritin trimerization domain (foldon) at the C terminus (Wrapp et al., 2020) followed by an HRV3C cleavage site, His8 and Twin-Streptactin purification tags. Additionally, the furin cleavage sites between S1 and S2 were mutated; for HKU1 and OC43, RRKRR was replaced by GGSGG; for MERS-CoV, RSVR was replaced with ASVG; for SARS-CoV, SHC014, WIV1, and RaTG13, SLLRST was replaced with SLLAST. DNA sequences encoding diverse beta-coronavirus S2P proteins were cloned into mammalian expression vector pVRC8400 and the proteins expressed by transient transfection of FreeStyle 293-F cells as previously described (Zhou et al., 2020b). Specifically, 1 mg of transfection grade plasmid DNA and 3 ml of Turbo293 transfection reagent (Speed BioSystems) each in 20ml Opti-MEM (Thermo Fisher Scientific), were pre-mixed and transfected into FreeStyle 293-F cells. Transfected cells were grown for 6 days at 37 ºC, and the supernatant was harvested by centrifugation to remove cell debris. Supernatants were sterile-filtered, and the spike trimers were purified by nickel affinity chromatography using cOmplete His-Tag purification resin. The resin was washed with 50 mM Tris, 400 mM NaCl, 10 mM and 25 mM Imidazole pH 8.0 buffer. The proteins were eluted in 50 mM Tris pH 8.0, 400 mM NaCl, 300 mM Imidazole. Protein fractions were pooled and further purified by size-exclusion chromatography (SEC) using a Superdex 200 16/600 column (Cytiva) in PBS. Fractions corresponding to trimeric spike proteins were pooled, concentrated to 1 mg/ml.

#### Production of antibodies

Antibody heavy and light variable region gene sequences were synthesized (Gene Universal Inc, Newark DE) and subcloned into corresponding pVRC8400 vectors (https://www.addgene.org). The resulting plasmids of heavy and light chain pairs were co-transfected in Expi293F cells (Thermo Fisher) using Turbo293 transfection reagent (Speed BioSystems) as described previously (Wu et al., 2011). On day 6 post transfection, the culture supernatants were harvested, sterile filtered and loaded onto a protein A column. The columns were washed with PBS, and IgG proteins were eluted with a low pH IgG elution buffer (Pierce) and immediately neutralized with 1M Tris-HCl pH 8.0. Purified IgGs were subsequently dialyzed twice against PBS pH 7.4 using 10 kD dialysis cartridges (Pierce) and used for measurements.

#### Full-length spike constructs

Codon optimized cDNAs encoding full-length spike from SARS-CoV-2 (GenBank ID: QHD43416.1) were synthesized, cloned into the mammalian expression vector VRC8400 (Barouch et al., 2005) and confirmed by sequencing. Spike containing D614G amino acid change was generated using the wildtype spike sequence. Other variants containing single or multiple amino-acid changes in the spike gene from wildtype or D614G were made by mutagenesis using QuickChange lightning Multi Site-Directed Mutagenesis Kit (cat # 210515, Agilent) or via synthesis and cloning (Genscript). The spike variants tested are B.1.1.7 (H69del-V70del-Y144del-N501Y-A570D-D614G-P681H-T716I-S982A-D1118H), B.1.351 (L18F-D80A-D215G-(L242-244)del-R246I-K417N-E484K-N501Y-A701V), P.1 (L18F-T20N-P26S-D138Y-R190S-K417T-E484K-N501Y-D614G-H655Y-T1027I-V1176F), B.1.617.2 (T19R, G142D, del156-157, R158G, L452R, T478K, D614G, P681R, D950N), AY.1 (T19R, T95I, G142D, E156del, F157del, R158G, W258L, K417N, L452R, T478K, D614G, P681R, D950N), B.1.621 (T95I, insert144T, Y144S, Y145N, R346K, E484K, N501Y, D614G, P681H, D950N) and B.1.1.529 (A67V, H69del, V70del, T95I, G142D, V143del, Y144del, Y145del, N211del,L212I, ins214EPE, G339D, S371L, S373P, S375F, K417N, N440K, G446S, S477N, T478K, E484A, Q493R, G496S, Q498R, N501Y, Y505H, T547K, D614G, H655Y, N679K, P681H, N764K, D796Y, N856K, Q954H, N969K, L981F). The spike genes from SARS-CoV-2 related CoVs (RaTG13_GenBank: QHR63300.2; Pangolin GD_GenBank: QIA48632.1; Pangolin_GX-P2V_ QIQ54048.1), SARS-CoV and related CoVs (SARS-CoV Urbani_ GenBank: AAP13441.1; Frankfurt1_GenBank: BAE93401.1; Civet SARS CoV 007/2004 S_ GenBank: AAU04646.1; WIV1_GenBank: KF367457; SHC014_GenBank: KC881005), MERS-CoV EMC_ GenBank: AFS88936), hCoV-HKU1 (GenBank: AAT98580.1), hCoV-OC43 (GenBank: AAT84354.1), hCoV-NL63 (GenBank: Q6Q1S2.1) and hCoV-229E (GenBank: AOG74783.1) were synthesized (Genscript). These full-length spike plasmids were used for pseudovirus production and for cell surface binding assays.

#### Analysis of WS6 binding to cell surface expressed spike protein

Expi-293 cells were transiently transfected with plasmids encoding full-length spike proteins of coronaviruses using Turbo293 transfection reagent (Speed BioSystems) following manufacturer’ s protocol. After 40 hours, cells were harvested and incubated with monoclonal antibodies (1 μg/ml) for 30 minutes. Cells were washed and incubated with an allophycocyanin conjugated anti-human IgG (709-136-149, Jackson Immunoresearch Laboratories) for another 30 minutes, then washed and fixed with 1% paraformaldehyde (15712-S, Electron Microscopy Sciences). Flow cytometry data were acquired in a BD LSRFortessa X-50 flow cytometer (BD biosciences) and analyzed using Flowjo (BD biosciences).

#### Pseudovirus neutralization assay

Spike-containing lentiviral pseudovirions were produced by co-transfection of packaging plasmid pCMVdR8.2, transducing plasmid pHR’ CMV-Luc, a TMPRSS2 plasmid and S plasmids from human and animal coronaviruses (SARS-CoV-2 variants, SARS-CoV, MERS-CoV and SARS-CoV-2, SARS-CoV related coronaviruses) into 293T cells using Lipofectamine 3000 transfection reagent (L3000-001, ThermoFisher Scientific, Asheville, NC) (Naldini et al., 1996). 293T-ACE2 cells (provided by Dr. Michael Farzan) or 293 flpin-TMPRSS2-ACE2 cells (made at the VRC) were plated into 96-well white/black Isoplates (PerkinElmer, Waltham, MA) at 75,00 cells per well the day before infection of pseudovirus. Serial dilutions of mAbs were mixed with titrated pseudovirus, incubated for 45 minutes at 37°C and added to cells in triplicate. Following 2 h of incubation, wells were replenished with 150 ml of fresh media. Cells were lysed 72 h later, and luciferase activity was measured with Microbeta (Perking Elmer). Percent neutralization and neutralization IC50s, IC80s were calculated using GraphPad Prism 8.0.2.

#### Neutralization by fusion inhibition (post attachment inhibition)

BHK21-ACE2 or 293 flpin-TMPRSS2-ACE2 cells were placed on ice for one hour before incubating with SARS-CoV-2 spike pseudotyped lentivirus on ice for another hour to allow the virus to attach to ACE2. After 3 times of wash the cells were then incubated with mAbs on ice for additional one hour before returned to the incubator. Cells were lysed 72 h later, and luciferase activity was measured with Microbeta (Perking Elmer). Percent neutralization and neutralization IC50s, IC80s were calculated using GraphPad Prism 8.0.2.

#### Biolayer interferometry analysis of antibody binding affinity

Binding kinetics of WS6 IgG and related S2-stem helix antibodies to diverse CoV 2P-stabilized spike proteins (S2Ps) were measured by biolayer interferometry using an Octet HTX instrument (Sartorius). All assays were performed in tilted 384-well plates (Geiger Bio-one) in HBS-EP+ buffer with agitation set to 1000 rpm at 30ºC. The final volume for all solutions was 60-80 μl/well. Prior to running the assays, Anti Mouse Fc (AMC) biosensor tips were equilibrated in PBS for ∼5 minutes. WS6 antibody IgG at 10 μg/ml were loaded onto AMC biosensors for 60s. Biosensors were then equilibrated in HBS-EP+ buffer for 60s prior to measuring association with diverse beta-CoV S2P trimers in solution (0.0019 μM to 0.5μM) for 300s; trimer proteins were then allowed to dissociate for 600s. Parallel correction to subtract systematic baseline drift was carried out by subtracting the measurements recorded for a sensor loaded with antibody incubated in HBS-EP+ buffer. Data analysis and curve fitting were carried out using Octet analysis software, version 12.0 (Sartorius). Experimental data were fitted using a 1:1 interaction to determine apparent dissociation constants and k_on_ values.

#### Crystallization and structure determination

The N-terminal acetylated, C-terminal biotin-pegylated, 10-residue SARS-CoV-2 S2 peptide FKEELDKYFK (GenScript) was dissolved in 100% DMSO at 10 mg/mL and then diluted with PBS to 1 mg/mL. Fab WS6 was incubated with the peptide in a 1:2 molar ratio for 5 minutes to form complex. The Fab-peptide complex was concentrated to 10-15 mg/mL and screened for crystallization using 576 conditions from Hampton screen, Precipitant Synergy screen, and QIAGEN Wizard screen with a Mosquito robot mixing 100 nL reservoir solution and 100 nL protein solution per drop. Crystals were obtained from the solution containing 2% v/v PEG 400, 2 M (NH_4_)_2_SO_4_, 100 mM sodium acetate pH 5.5. Crystal drops were set up in 15-well plates using 0.5 μL WS6-peptide complex and 0.5 μL of 80-100% concentration of the aforementioned crystallization solution. Crystals were cryoprotected in the aforementioned solution supplemented with 50% additional precipitant and 15% v/v (2R,3R)-2,3-Butanediol and flash frozen in liquid nitrogen.

X-ray diffraction data were collected at the Advanced Photon Source (beamline SER-CAT ID22 and BM22). The diffraction data were indexed, integrated, and scaled with the HKL2000 package (Otwinowski and Minor, 1997). The structures were determined by molecular replacement with Phaser (McCoy et al., 2007) in the CCP4 Program Suite (Collaborative Computational Project, 1994). Further refinement was carried out with PHENIX (Adams et al., 2010), starting with torsion-angle simulated annealing with slow cooling. Iterative manual model building was carried out with COOT (Emsley and Cowtan, 2004) with maps generated from combinations of standard positional, individual B-factor and TLS refinement algorithms. X-ray data and refinement statistics are summarized in Table S2.

#### Antibody germline assignment

Germline gene of heavy and light chain were assigned by querying nucleotide sequences on IGBLAST server. B6 heavy and light chain germline gene were assigned by submitting protein sequence. The V gene identity was reported as the identity to the germline nucleotide sequences.

#### Antibody frequency calculation

The mouse Ig deep sequencing samples were prepared and used to predict probability of generation of specific protein sequences on CDR3. Human Ig deep sequencing samples were downloaded from Bioproject PRJNA511481. The Ig sequences were sieved for IGOR to inference the model (Marcou et al., 2018). The inferenced models were used by OLGA to predict probability of having specific CDR3 sequences (Sethna et al., 2019). Signature of CV3-25 class antibody was defined as heavy chain IGHV5-51 and light chain IGKV1-12 germline gene, the R94, 96PQYC, G100b, C100d, and W100g on 18 amino acids CDR H3. The frequency of S2P6 class antibody was calculated by defining signature as IGHV1-46 germline gene with 97PKG in 11 amino acids CDR H3, and IGKV3-20 with Y91, 93SSPP and 96F on 11 amino acids CDR L3. The frequency of cc40.8 class antibody was calculated by masking IGHV3-23 with 94ITMA, 101V and 102V on 8 amino acids CDR H3 and IGLV3-10 with T91, 94SGN, and A96 on 11 amino acids CDR L3. The frequency of WS6 class antibody calculated by using mouse IGHV1-5 gene with 95TGS on 7 amino acids CDR H3, and IGKV4-61 gene with 91Y, 94Y and 96P on 9 amino acids CDR L3. B6 and IgG22 used identical heavy chain germline gene, although the light chain used different germline genes, it shared similar contact residues. B6/IgG22 class frequency was calculated by IGHV1-19 germline gene with CX{5,10}RX{4,6}W signature on CDR H3 amino acids, and IGKV8-27 /IGKV1-99 germline genes with 92[LN] and 95[FY] on 8 amino acids length CDR L3.

#### Phylogenetic tree

Alpha-coronavirus spike sequences were downloaded using accession codes (NL63:Q6Q1S2.1, 229E:AOG74783.1, Feline:AAY32596.1, Mink:ADI80513.1). Beta-coronavirus spike sequences were downloaded from NCBI server using accession codes listed in Full-length spike constructs. ClustalW was used to calculate neighbor joining (NJ) tree (Larkin et al., 2007), and Dendroscope was used to plot the Neighbor Joining tree (Albrecht et al., 2012).

### QUANTIFICATION AND STATISTICAL ANALYSIS

The BLI data were analyzed and plotted using GraphPad Prism. Crystal diffraction data and structural refinement statistics were analyzed with HKL2000, Phenix, and Molprobity. Statistical details of experiments are described in Method Details or figure legends.

