## Supplemental for "Vaccine-elicited murine antibody WS6 neutralizes diverse beta-coronaviruses by recognizing a helical stem supersite of vulnerability"

Figure S1

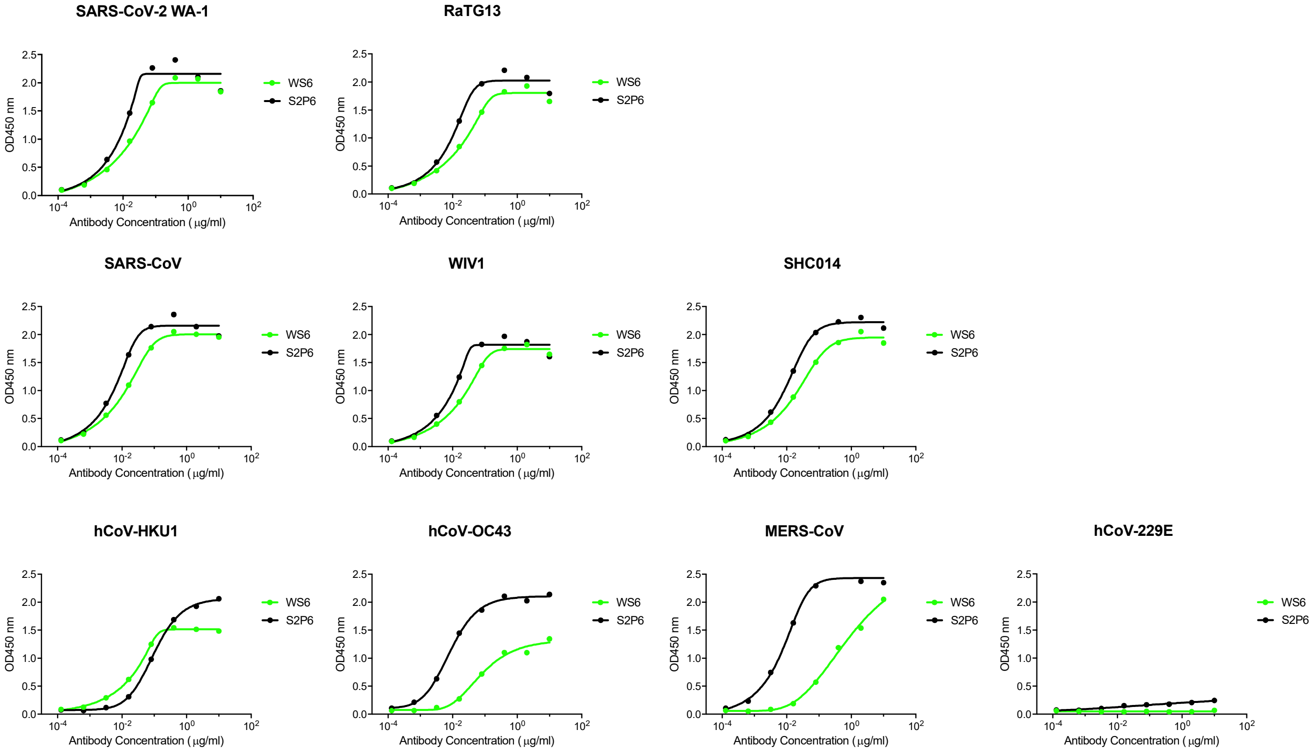


**Figure S1. Binding of WS6 to diverse CoV spike proteins by ELISA compared to S2P6, related to Figures 1 and 2.** ELISA binding curves of WS6 are shown in green and those of S2P6 in black.

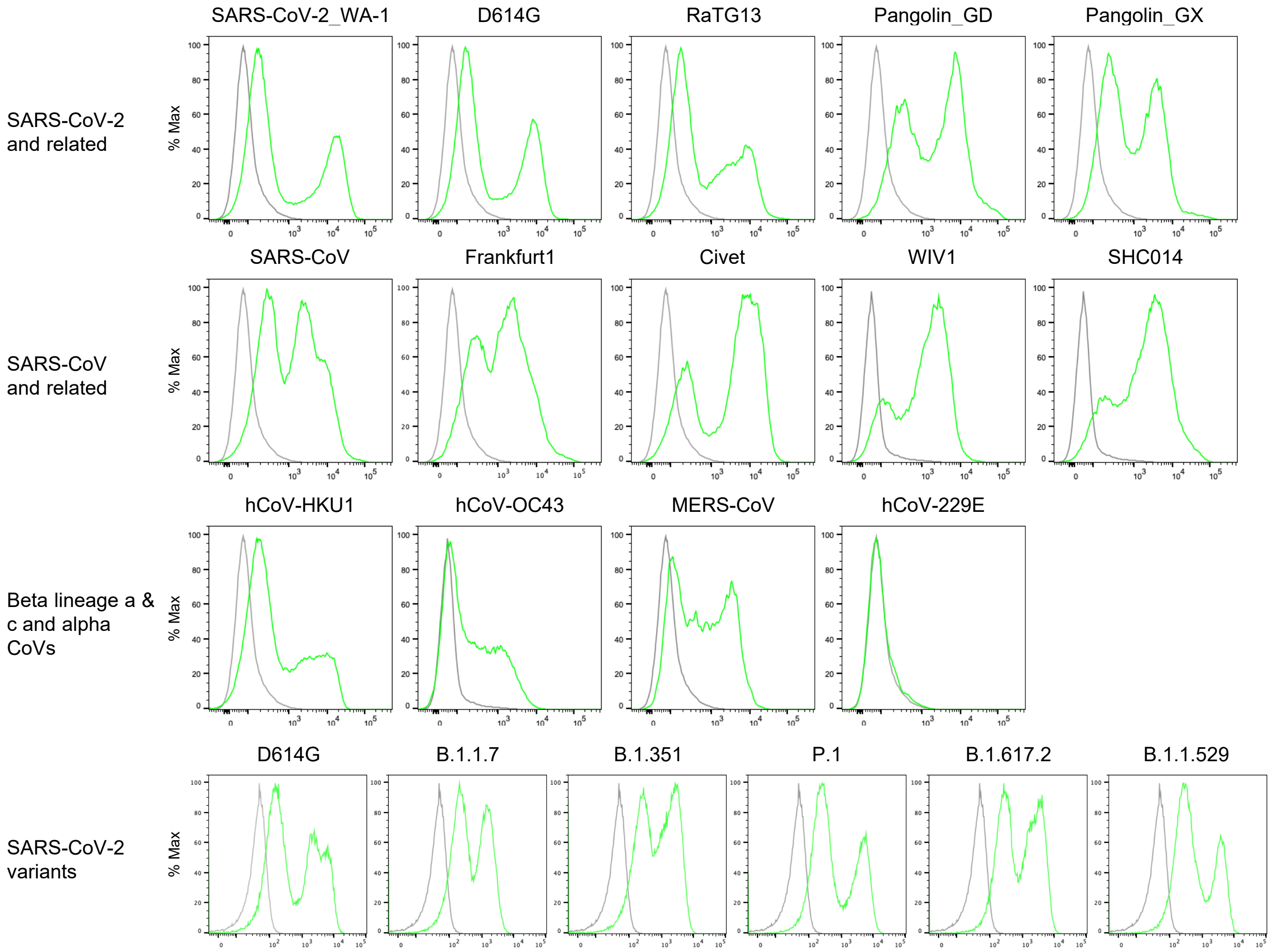


**Figure S2. WS6 binding to cell-surface expressed coronavirus spikes, related to Figure 2.**
Spike proteins were expressed on the surface of expi293 cells, and antibody binding was measured using flow cytometry. WS6 binding (green line) to cells transfected with the indicated coronavirus spike is compared to binding to untransfected cells (grey line).

Figure S3

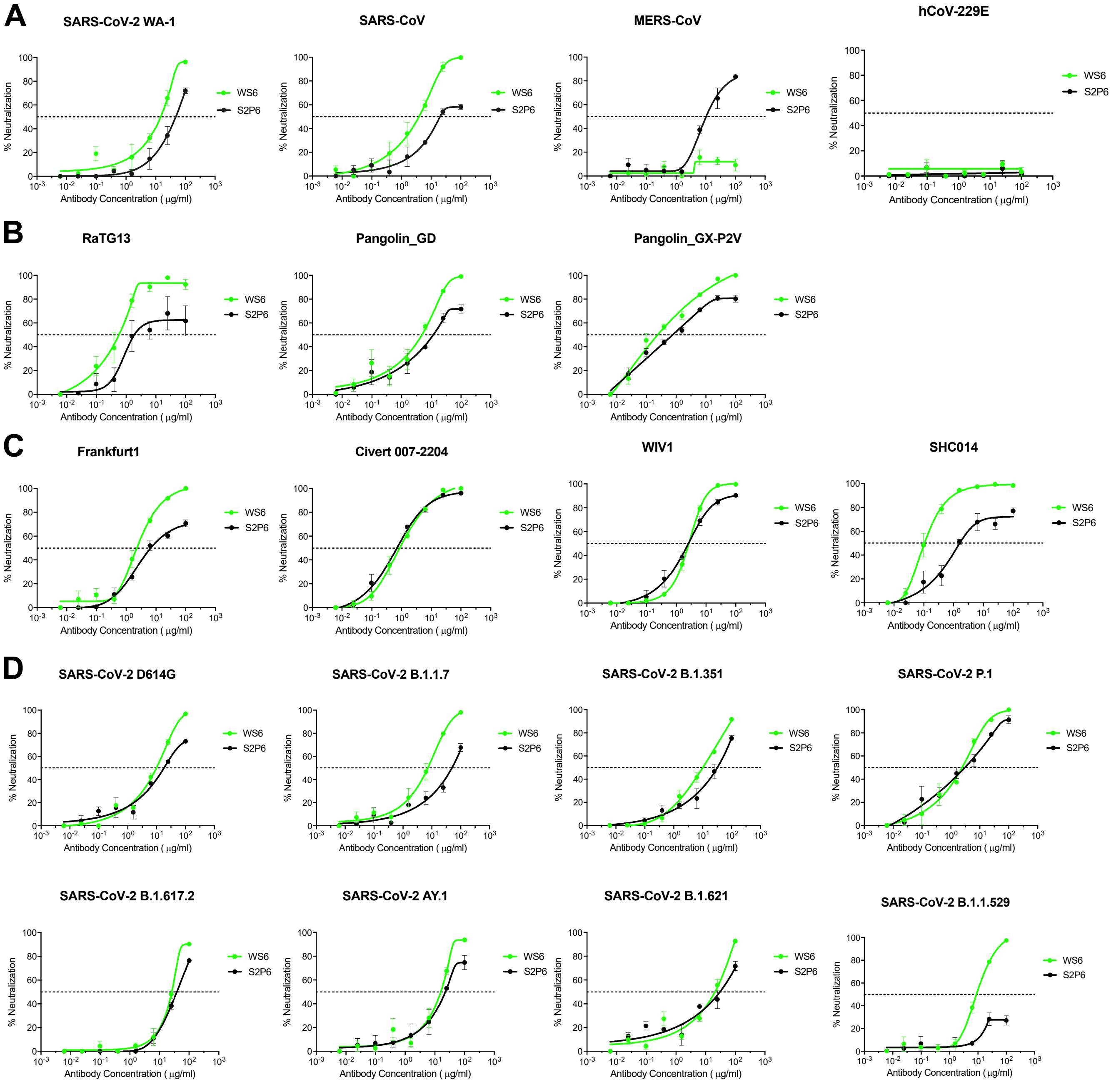


**Figure S3. Neutralization of WS6 against diverse coronaviruses, related to Figure 2.**
Neutralization curves are showing using CoV spike pseudotyped lentivirus to test neutralization capacity of WS6 compared to S2P6. Neutralization was tested on HEK293-TMPRSS2-ACE2 stable cells for SARS-CoV-2, SARS-CoV and related CoVs and Huh7.5 cells for MERS-CoV. (**A**) WS6 neutralizes SARS-CoV-2, SARS-CoV, but not MERS-CoV or hCoV-229E. (**B**) WS6 neutralizes SARS-CoV-2 related coronaviruses. (**C**) WS6 neutralizes SARS-CoV related coronaviruses. (**D**) WS6 neutralizes SARS-CoV-2 variants.

**A**

```
                 1138      1148    1156                                                      1208
SARS-2 (1273)    YDPLQPELDS-------FKEELDKYFKNHTSPDVDLG-------DISGINASVVNIQKEIDRLNEVAKNLNESLIDLQELGKYEQ
RaTG13(1269)     YDPLQPELDS-------FKEELDKYFKNHTSPDVDLG-------DISGINASVVNIQKEIDRLNEVAKNLNESLIDLQELGKYEQ
Pangolin_GD(1267) YDPLQPELDS-------FKEELDKYFKNHTSPDVDLG-------DISGINASVVNIQKEIDRLNEVAKNLNESLIDLQELGKYEQ
Pangolin_GX(1269) YDPLQPELDS-------FKEELDKYFKNHTSPDVDLG-------DISGINASVVNIQKEIDRLNEVAKNLNESLIDLQELGKYEQ
SARS(1255)       YDPLQPELDS-------FKEELDKYFKNHTSPDVDLG-------DISGINASVVNIQKEIDRLNEVAKNLNESLIDLQELGKYEQ
WIV1(1256)       YDPLQPELDS-------FKEELDKYFKNHTSPDVDLG-------DISGINASVVNIQKEINRLNEVAKNLNESLIDLQELGKYEQ
SHC014(1256)     YDPLQPELDS-------FKEELDKYFKNHTSPDVDLG-------DISGINASVVNIQKEIDRLNEVAKNLNESLIDLQELGKYEQ
MERS(1353)       STNLPPPLLGNSTGI-DFQDELDEFFKNVSTSIPNFG-------SLTQINTTLLDLTYEMLSLQQVVKALNESYIDLKELGNYTY
OC43(1353)       LNTSTPNLPD-------FREELDQWFKNQTSVAPDL--------SLDYINVTFLDLQVEMNRLQEAIKVLNQSYINLKDIGTYEY
HKU1(1356)       LNNSIPNLSD-------FEAELSLWFKNHTSIAPNLT-------FNSHINATFLDLYYEMNVIQESIKSLNSSFINLKEIGTYEM
NL63(1356)       LHTVIPDYVDVNKTLQEFAQNLPKYVKPNFDLTPFNLTYLNLSSELKQLEAKTASLFQTTVELQGLIDQINSTYVDLKLLNRFEN
229E(1171)       LQTIVPEYIDVNKTLQELSYKLPNYTVPDLVVEQYNQTILNLTSEISTLENKSAELNYTVQKLQTLIDNINSTLVDLKWLNRVET
```

**B** Germline gene usage of antibodies targeting S2-helical region

| Antibody | VH | VH Identity (%) | CDRH3 | VL | VL Identity (%) | CDRL3 | Ref |
|---|---|---|---|---|---|---|---|
| CV3-25 | IGHV5-51 | 97.6 | CARLPQYCSNGVCQRWFDPW | VK1-12 | 97.5 | CQQGNSFPYTF | (Li et al., 2022) |
| S2P6 | VH1-46 | 96.5 | CARGSPKGAFDYW | VK3-20 | 97.5 | CQQYGSSPPRFTF | (Pinto et al., 2021) |
| CC40.8 | VH3-23 | 93.8 | CAITMAPVVW | VL3-10 | 96.2 | CYSTDSSGNHAVF | (Song et al., 2021) |
| WS6 | VH1-5 | 92.9 | CTRTGSY-FDYW | VK4-61 | 97.9 | CQQYQSYPPTF | This study |
| B6 | VH1-19 | - | CARQLGRGNGLDYW | VK8-27 | - | CHQYLSSYTF | (Sauer et al., 2021) |
| IgG22 | VH1-19 | 95.2 | CTRVRGNDYHGRAMDYW | VK1-99 | 98.6 | CFQSNYLFTF | (Hsieh et al., 2021) |

Blue: human antibody
Green: mouse antibody

**C** Heavy and light chain alignment

```
Heavy Chain
         <-------------FR1------------><CDR><-----FR2----><-----CDR2------><--------------FR3-------------><-----CDR3-----><-----FR4->
CV3-25   EVQLVESGAEVKKPGESLKISCKGSGYTFTRYWIGWVRQMPGKGLEWMGIIYPGDSDTRYSPSFQGHVTISADKSISTAYLQWNSLKASDTAMYYCARLPQYCSNGVCQRWFDPWGQGTLVTVSS
S2P6     EVQLVQSGAEVKKPGASVKVSCKASGYTFTSQYMHWVRQAPGQGLEWIGIINPSGVHTSYAQKFQGRVTLTRDTSTSTLYMELSSLRSEDTAVYYCAR-------GSPKGAFDYWGQGTLVTVSS
CC40.8   EVQLLESGGGLVQPGGSLRLSCAASGFTFSSYVMTWARQAPGKGLEWVSAI-SGTGYTYYADSVKGRFTVSRDNSKNTLFLQMSSLRAEDTAVYYCAITM----------APVVWGQGTTVTVSS
WS6      EVQFQQSGTVLARPGASVKMSCKASGYTFTNYWIHWVKQRPGQGLEYIGGTYPGNGDTTYNQKFKGKAKVTAVTPTSTAYMDLSSLTNEDSAVYYCTRT---------GSYFDYWGQGTTLTVSS
B6       EVQLQQSGPVLVKPGASVRMSCKASGYTITDYYLNWVKQSHGKSLEWLGVLNPYSGGSLYSQTFKGKATLTVDRSSSTAYLELNSLTSEDSAVYYCARQL------GRGNGLDYWGQGTSVTVSS
IgG22    EVQLQQPGPVLVKPGASVRMSCKASGYRITDNFMNWVKQSHGKSLEWIGIINPYNGGTKYNQKFKGKATLTVDTSSSTAYMELNSLTSEDSAVYYCTRVRGN-DYHGR--AMDYWGQGTSVTVSS
Light Chain
         <-------FR1-----------><------CDR1-----><-----FR2-----><CDR2-><--------------FR3-------------><---CDR3--><--FR4--->
CV3-25   EIVLTQSPSSVSASVGDRVTITCRASQGI------SSWLAWYQQKPGKAPKLLIYAASSLQSGVPSRFSGSGSGTDFTLTISSLQPEDFATYYCQQGNSFP--YTFGQGTNLEIK
S2P6     EIVMMQSPGTLSLSPGERATLSCRASQSV-----RSNYLAWYQQKPGQAPRLLIYGASSRATGIPDRFSGSGSGTDFTLTISRLEPEDFAVYYCQQYGSSPPRFTFGPGTKVEIK
CC40.8   SYELTQPPS-VSVSPGQTARITCSGDALP------KRYAYWYQQKSGQAPILVIYEDKKRPSGIPERLSGSKSGTVATLTISGAQVEDEADYYCYSTDSSGNHAVFGGGTQLTVL
WS6      QIVLTQSPAIMSASPGEKVTISCSATSSV-------SYIYWYQQRPGSSPKPWIYRTSNLASGVPVRFSGSGSGTSYSLTISNMEAEDAATYYCQQYQSYPP--TFGAGTKLELK
B6       NIMMTQSPSSLAVSAGEKVTMSCKSSQSVLHSSDQKNYLAWYQQKPGQSPKLLIYWASTRESGVPDRFTGSGSGTDFTLTISSVQAEDLAVYFCHQYLSS---YTFGGGTKLEIK
IgG22    DVVLTQTPLSLPVNIGDQASISCKSTKSLLNR-DGFTFLDWYLQKPGQSPQLLIYLVSNRFSGVPDRFSGSGSGTDFTLKISRVEAEDLGVYYCFQSNYL---FTFGSGTKLEIK
```

**D** Frequency of antibodies targeting S2-helical region

| | | HC Frequency | LC Frequency | Class Frequency | Average Class Frequency |
|---|---|---|---|---|---|
| CV3-25 | HIP1 | 8.11E-10 | 5.46E-03 | 2.65E-12 | 1.90E-12 |
| | HIP2 | 4.58E-10 | 4.11E-03 | 1.13E-12 | |
| | HIP3 | 6.39E-10 | 5.02E-03 | 1.92E-12 | |
| S2P6 | HIP1 | 6.54E-09 | 2.82E-04 | 1.10E-12 | 1.71E-12 |
| | HIP2 | 5.81E-09 | 3.01E-04 | 1.05E-12 | |
| | HIP3 | 1.37E-08 | 3.63E-04 | 2.99E-12 | |
| CC40.8 | HIP1 | 4.09E-14 | 5.67E-07 | 9.27E-21 | 2.52E-20 |
| | HIP2 | 5.58E-14 | 1.68E-06 | 3.76E-20 | |
| | HIP3 | 6.57E-14 | 1.10E-06 | 2.88E-20 | |
| WS6 | Mouse1 | 3.57E-08 | 2.90E-05 | 9.83E-13 | 5.00E-13 |
| | Mouse4 | 2.60E-08 | 1.56E-06 | 3.84E-14 | |
| | Mouse5 | 7.11E-08 | 7.07E-06 | 4.78E-13 | |
| B6/IgG22 | Mouse1 | 1.38E-04 | 2.92E-04 | 3.83E-08 | 3.72E-08 |
| | Mouse4 | 1.39E-04 | 3.67E-04 | 4.84E-08 | |
| | Mouse5 | 2.38E-04 | 1.10E-04 | 2.48E-08 | |

**Figure S4. Sequences of conserved S2 stem region and antibodies that target this region, related to Figure 5.**
(**A**) S2-stem sequences of diverse coronaviruses. Aqua highlight for amino acids conserved on SARS/SARS2. Yellow/red font shown full conserve. Green highlight shown amino acids with similar physicochemical property. (**B**) Identified human and mouse antibodies targeted on coronavirus spike S2-Helix epitope. (**C**) Alignments of heavy and light chain sequences. Residues that contact the helix epitope are highlighted in cyan. (**D**) The frequencies of antibody targeting SP2 helical region calculated by software OLGA. See Methods for the signatures used to calculate frequency.

### Figure S5

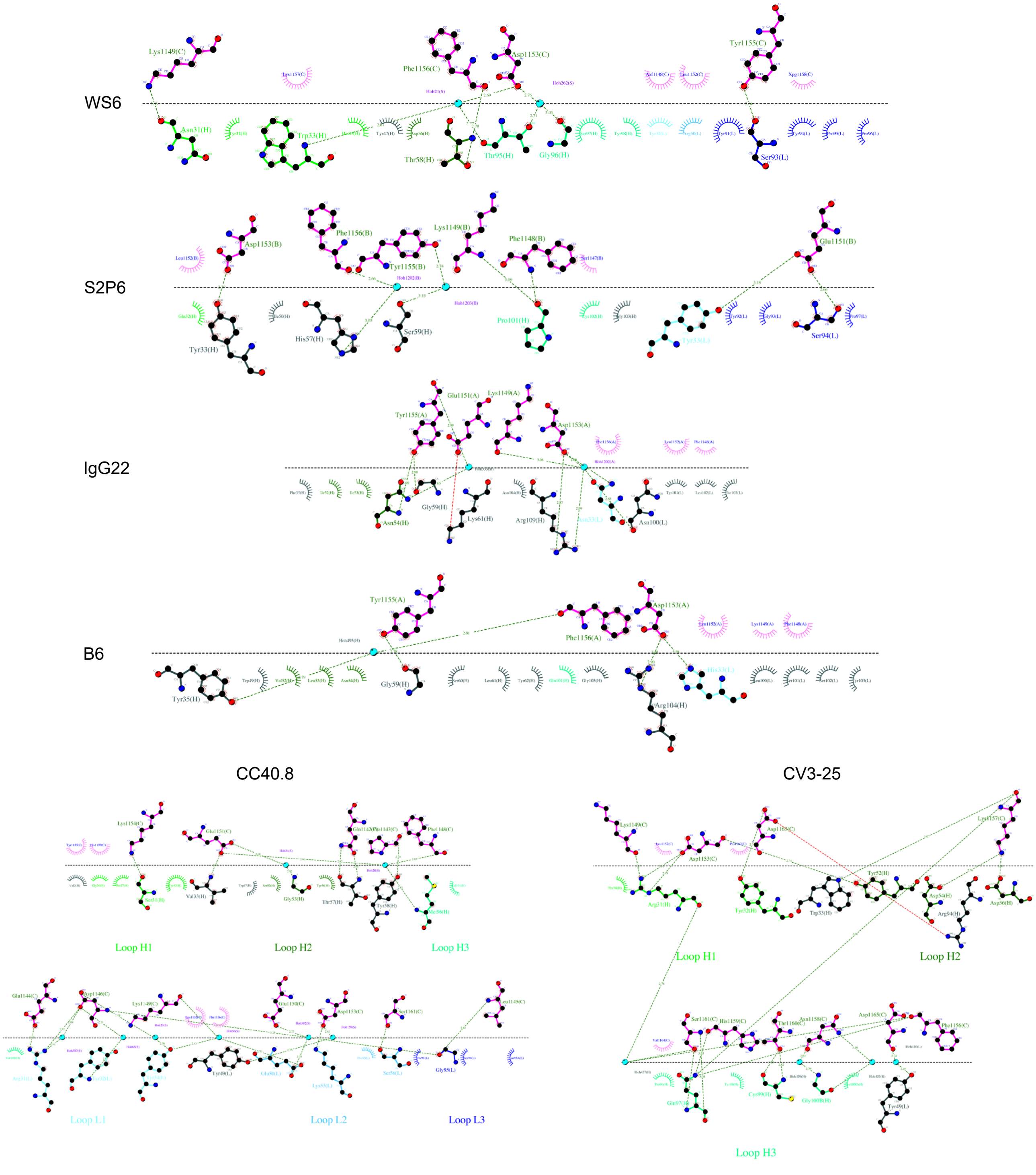


**Figure S5. Detailed interactions between antibodies and S2 peptide residues, related to Figure 5.**
Hydrophilic and hydrophilic interaction between antibody and S2 residues were plotted with LigPlot+ (Laskowski R A, Swindells M B (2011). LigPlot+: multiple ligand-protein interaction diagrams for drug discovery. J. Chem. Inf. Model., 51, 2778-2786. [PubMed id: 21919503]). Interface is depicted with a horizontal line between antibody and peptide residues. Hydrophilic interaction are shown as lines between atoms with distance labeled. Hydrophobic interactions are shown as eyelash symbols.

**Table S1. Pseudovirus neutralization of WS6 of diverse coronaviruses, related to Figures 1 and 2.**

| Pseudoviruses | WS6 | | S2P6 | |
|---|---|---|---|---|
| | IC50 | IC80 | IC50 | IC80 |
| WA-1* | 4.28 | 24.01 | 16.12 | 76.39 |
| D614G | 9.88 | 34.86 | 17.55 | >100 |
| B.1.1.7 | 6.31 | 26.81 | 49.25 | >100 |
| B.1.351 | 9.83 | 53.74 | 32.02 | >100 |
| P.1 | 2.46 | 10.45 | 3.06 | 28.39 |
| B.1.617.2 | 26.52 | 69.85 | 38.90 | >100 |
| Delta+ | 15.59 | 32.37 | 22.46 | >100 |
| B.1.621 | 20.26 | 63.82 | 31.05 | >100 |
| B.1.529 | 3.43 | 35.76 | >100 | >100 |
| RaTG13 | 0.52 | 1.79 | 1.74 | >100 |
| Pangolin_GD | 4.91 | 18.34 | 11.71 | >100 |
| Pangolin_GX | 0.24 | 3.89 | 0.76 | 27.22 |
| SARS | 1.93 | 9.57 | 24.03 | >100 |
| Frankfurt 1 | 2.27 | 8.63 | 6.72 | >100 |
| Civet 007-2004 | 0.84 | 4.67 | 0.68 | 4.32 |
| WIV1 | 2.65 | 6.41 | 2.44 | 15.87 |
| SHC014 | 0.11 | 0.39 | 1.57 | >100 |
| MERS-CoV | >100 | >100 | 10.59 | >100 |
| hCoV-229E | >100 | >100 | >100 | >100 |

| ug/ml |
|---|
| 0.1-1 |
| 1-10 |
| 10-100 |
| >100 |

* The titers shown here reflect neutralization tested on HEK293-TMPRSS2-ACE2 stable cells and differ from those in Figure 1D, which were tested on 293T-ACE2 cells.

**Table S2. Crystal diffraction data and structure refinement statistics, related to Figure 3.**

| | |
|---|---|
| Wavelength (Å) | 1.0000 |
| Resolution range (Å) | 40.23 - 2.02 (2.092 - 2.02) |
| Space group | C 1 2 1 |
| Unit cell (a, b, c, $\alpha$, $\beta$, $\gamma$) | 155.1, 64.5, 138.0, 90.0, 117.3, 90.0 |
| Total reflections | 262622 |
| Unique reflections* | 75855 (6642) |
| Multiplicity | 3.5 (2.9) |
| Completeness (%) | 94.81 (81.69) |
| Mean I/sigma(I) | 10.5 (1.1) |
| Wilson B-factor | 27.88 |
| R-merge | 0.101 (0.943) |
| R-pim | 0.083(0.513) |
| CC1/2 | 0.995 (0.765) |
| Reflections used in refinement | 75540 (6475) |
| Reflections used for R-free | 1989 (170) |
| R-work | 0.1718 (0.2427) |
| R-free | 0.2133 (0.2848) |
| Number of non-hydrogen atoms | 7587 |
| macromolecules | 6696 |
| ligands | 176 |
| solvent | 735 |
| Protein residues | 871 |
| RMS(bonds, Å) | 0.008 |
| RMS(angles, °) | 0.91 |
| Ramachandran favored (%) | 97.78 |
| Ramachandran allowed (%) | 2.22 |
| Ramachandran outliers (%) | 0.00 |
| Rotamer outliers (%) | 0.92 |
| Clashscore | 4.58 |
| Average B-factor (Å$^2$) | 32.90 |
| macromolecules | 31.84 |
| ligands | 49.90 |
| solvent | 38.92 |

* Statistics for the highest-resolution shell are shown in parentheses.

**Table S3. WS6-peptide binding interface analysis, related to Figure 3.**

The crystal structure of WS6 in complex with the stem-helix peptide was analyzed by PISA (https://www.ebi.ac.uk/msd-srv/prot_int/cgi-bin/piserver). ASA, accessible surface area in Å$^2$; BSA, buried surface area in Å$^2$; ΔiG, solvation free energy gain upon formation of the interface in kcal/M. Bars of BSA indicates buried area percentage, one bar per 10%. Atoms with superscript "H" are involved in interface hydrogen bonds.

<table>
<tr><th>WS6</th><th>ASA</th><th>BSA</th><th>ΔiG</th><th>CDR-BSA</th><th>Peptide</th><th>ASA</th><th>BSA</th><th>ΔiG</th></tr>
<tr><td colspan="9">Heavy chain interface</td></tr>
<tr><td>H:Asn31<br>[ O ]$^{H}$ [ CB ]</td><td>87.76</td><td>17.02 ||</td><td>-0.18</td><td>CDR H1<br>126.61</td><td>C:Phe1148<br>[ C ][ O ][ CB ]</td><td>380.59</td><td>39.54||</td><td>0.00</td></tr>
<tr><td>H:Tyr32<br>[ CA ][ CD2][ CE2][ CZ ][ OH ]</td><td>71.35</td><td>19.09 |||</td><td>0.29</td><td></td><td>C:LYS1149<br>[ N ][ CA ][ CB ][ CG ][ CD ][ CE ][ NZ ]$^{H}$</td><td>182.72</td><td>103.04||||||</td><td>0.41</td></tr>
<tr><td>H:Trp33<br>[ N ][ CB ][ CG ][ CD1][ CD2][ CE2][ CE3][ NE1]$^{H}$<br>[ CZ2][ CZ3][ CH2]</td><td>81.33</td><td>75.56 ||||||||||</td><td>0.67</td><td></td><td>C:Leu1152<br>[ CB ][ CG ][ CD2]</td><td>100.74</td><td>55.93||||||</td><td>0.89</td></tr>
<tr><td>H:His35<br>[ CE1]</td><td>14.94</td><td>14.94 |||||||||||</td><td>0.24</td><td></td><td>C:Asp1153<br>[ CA ][ O ]H [ CB ][ CG ][ OD1][ OD2]</td><td>68.05</td><td>41.86|||||||</td><td>-0.16</td></tr>
<tr><td>H:Tyr47<br>[ CD1][ CD2][ CE1][ CE2][ CZ ][ OH ]</td><td>74.50</td><td>14.78 ||</td><td>0.13</td><td>CDR H2<br>87.62</td><td>C:PHE1156<br>[ CA ][ C ][ O ]$^{H}$ [ CB ][ CG ][ CD1] [ CD2] [ CE1][ CE2][ CZ ]</td><td>152.52</td><td>105.56|||||||</td><td>1.16</td></tr>
<tr><td>H:Tyr52<br>[ CE1][ CZ ][ OH ]</td><td>59.05</td><td>7.43 ||</td><td>0.10</td><td></td><td>C:Lys1157<br>[ N ][ CA ][ C ][ O ][ CB ][ CG ][ CD ][ CE ]</td><td>210.66</td><td>57.36|||</td><td>0.58</td></tr>
<tr><td>H:Asn55<br>[ ND2]</td><td>101.01</td><td>2.18 |</td><td>-0.02</td><td></td><td></td><td></td><td></td><td></td></tr>
<tr><td>H:Asp57<br>[ CB ][ CG ][ OD1][ OD2]</td><td>73.89</td><td>29.15 ||||</td><td>-0.17</td><td></td><td></td><td></td><td></td><td></td></tr>
<tr><td>H:Thr59<br>[ CB ][ CG2][ OG1]$^{H}$</td><td>78.40</td><td>34.08 |||||</td><td>0.28</td><td></td><td></td><td></td><td></td><td></td></tr>
<tr><td>H:Thr99<br>[ O ][ CB ][ CG2][ OG1]</td><td>27.78</td><td>22.48 |||||||||</td><td>0.13</td><td>CDR H3<br>145.3</td><td></td><td></td><td></td><td></td></tr>
<tr><td>H:Gly100<br>[ CA ][ C ][ O ]</td><td>44.61</td><td>42.18 |||||||||</td><td>-0.15</td><td></td><td></td><td></td><td></td><td></td></tr>
<tr><td>H:Ser101<br>[ N ][ CA ][ C ][ O ][ CB ][ OG ]</td><td>107.81</td><td>68.21 |||||||</td><td>0.93</td><td></td><td></td><td></td><td></td><td></td></tr>
<tr><td>H:Tyr102<br>[ CA ]</td><td>151.61</td><td>4.68 |</td><td>0.07</td><td></td><td></td><td></td><td></td><td></td></tr>
<tr><td>H:Phe103<br>[ CZ ]</td><td>82.75</td><td>7.75 |</td><td>0.12</td><td></td><td></td><td></td><td></td><td></td></tr>
<tr><td colspan="9">Light chain interface</td></tr>
<tr><td>L:Tyr31<br>[ CE2][ CE2][ OH ]</td><td>106.29</td><td>23.26 ||</td><td>0.00</td><td>CDR L1<br>23.26</td><td>C:Phe1148<br>[ CB ][ CG ][ CD1][ CD2][ CE1][ CE2][ CZ ]</td><td>380.59</td><td>75.6||</td><td>0.00</td></tr>
<tr><td>L:Arg49<br>[ CD ][ NE ][ CZ ][ NH1][ NH2]</td><td>112.09</td><td>54.24 |||||</td><td>-0.42</td><td>CDR L2<br>54.24</td><td>C:Lys1149<br>[ N ]</td><td>182.72</td><td>0.15|</td><td>-0.00</td></tr>
<tr><td>L:Tyr90<br>[CA][ O ][CB][CG][CD1][CD2][CE1][CE2][ CZ ][ OH ]</td><td>122.46</td><td>72.86 ||||||</td><td>0.63</td><td>CDR L3<br>151.58</td><td>C:Glu1151<br>[ OE1]</td><td>146.85</td><td>6.74|</td><td>-0.13</td></tr>
<tr><td>L:Gln91<br>[ O ]</td><td>91.56</td><td>1.71 |</td><td>-0.02</td><td></td><td>C:Leu1152<br>[ CD1][ CD2]</td><td>100.74</td><td>39.55||||</td><td>0.63</td></tr>
<tr><td>L:Ser92<br>[ CA ][ C ][ O ]$^{H}$</td><td>52.85</td><td>4.77 |</td><td>0.03</td><td></td><td>C:Tyr1155<br>[ O ][ CD2][ CE1][ CE2][ CZ ][ OH ]$^{H}$</td><td>164.25</td><td>112.17|||||||</td><td>0.36</td></tr>
<tr><td>L:Tyr93<br>[ N ][ CA ][ CB ][ CD2][ CE2][ CZ ][ OH ]</td><td>196.27</td><td>54.06 |||</td><td>0.80</td><td></td><td>C:Phe1156<br>[ CA ][ O ][ CD1][ CE1][ CZ ]</td><td>152.52</td><td>38.32|||</td><td>0.48</td></tr>
<tr><td>L:Pro95<br>[ CG ][ CD ]</td><td>57.14</td><td>18.18 ||||</td><td>0.29</td><td></td><td></td><td></td><td></td><td></td></tr>
</table>
